# Transcriptional Regulation and Mechanism of SigN (ZpdN), a pBS32 encoded Sigma Factor

**DOI:** 10.1101/624585

**Authors:** Aisha T. Burton, Aaron DeLoughery, Gene-Wei Li, Daniel B. Kearns

## Abstract

Laboratory strains of *Bacillus subtilis* encodes as many as 16 alternative sigma factors, each dedicated to expressing a unique regulon such as those involved in stress resistance, sporulation, and motility. The ancestral strain of *B. subtilis* also encodes an additional sigma factor homolog, ZpdN, not found in lab strains due to it being encoded on the large, low copy number plasmid pBS32 that was lost during domestication. DNA damage triggers pBS32 hyper-replication and cell death in a manner that depends on ZpdN but how ZpdN mediates these effects was unknown. Here we show that ZpdN is a bona fide sigma factor that can direct RNA polymerase to transcribe ZpdN-dependent genes and we rename ZpdN to SigN accordingly. Rend-seq analysis was used to determine the SigN regulon on pBS32, and the 5’ ends of transcripts were used to predict the SigN consensus sequence. Finally, we characterize the regulation of SigN itself, and show that it is transcribed by at least three promoters: *P_sigN1_*, a strong SigA-dependent LexA-repressed promoter, *P_sigN2_*, a weak SigA-dependent constitutive promoter, and *P_sigN3_*, a SigN-dependent promoter. Thus, in response to DNA damage LexA is derepressed, SigN is expressed and then experiences positive feedback. How cells die in a pBS32-dependent manner remains unknown, but we predict that death is the product of expressing one or more genes in the SigN regulon.

**IMPORTANCE:** Sigma factors are utilized by bacteria to control and regulate gene expression. Extra cytoplasmic function sigma factors are activated during times of stress to ensure the survival of the bacterium. Here, we report the presence of a sigma factor that is encoded on a plasmid that leads to cellular death after DNA damage.

## INTRODUCTION

Propagation and cultivation of bacteria in the laboratory has been shown to select for enhanced axenic growth and genetic tractability in a process called domestication. The model genetic bacterium *Bacillus subtilis* is an example of a commonly used domesticated bacterium as the laboratory strains differ substantially from the ancestor from which they were derived. For example, lab strains are defective for biofilm formation, reduced for motility, are auxotrophic for one or more amino acids, and are deficient in the ability to synthesize multiple antibiotics, a potent surfactant, and viscous slime layer production (1–5). While many traits were lost during the domestication of laboratory strains, one important trait was gained: high frequency uptake of extracellular DNA in a process called natural genetic competence. Later it was shown that increased genetic competence was also due to genetic loss, in this case due to the loss of the endogenous plasmid

pBS32 (6, 7). pBS32, is a large, 84 kb, low copy number plasmid that has a separate replication initiation protein and a high-fidelity plasmid partitioning system (6, 8–10). Moreover, pBS32 been shown to encode an inhibitor of competence for DNA uptake (ComI) (7) and an inhibitor of biofilm formation (RapP) that directly regulate cell physiology (11–13). In addition, approximately one third of the pBS32 sequence encodes a cryptic prophage like element, cell death is triggered in a pBS32-dependent manner following treatment with the DNA damaging agent, mitomycin C (MMC), and MMC often induces prophage conversion (7, 14–17). pBS32-dependent cell death upon mitomycin C treatment requires a plasmid-encoded sigma factor homolog, ZpdN and artificial ZpdN induction was shown to be sufficient to trigger cell death (17). How ZpdN is activated by the presence of DNA damage and the mechanism by which ZpdN promoted cell death was unknown.

Here we show that ZpdN functions as a *bona fide* sigma factor which directs RNA polymerase to transcribe a large regulon of genes encoded on pBS32. Based on our findings we rename ZpdN to SigN and propose a SigN-dependent consensus sequence for transcriptional activation. We show that SigN induction triggers immediate loss of cell viability, even as cells continue to grow and the cell culture increases in optical density. We characterize the *sigN* promoter region and find multiple promoters that activate its expression including a DNA damage-responsive LexA-repressed promoter and a separate promoter that governs autoactivation. Finally, the SigN regulon does not appear to include the pBS32 putative prophage region and thus cell death may be prophage independent. The gene or genes responsible for pBS32-mediated cell death remain unknown but we infer that they must reside within the plasmid expressed by SigN and RNA polymerase.

## MATERIALS AND METHODS

### Strains and growth conditions

*B. subtilis* strains were grown in lysogeny broth (LB) (10 g tryptone, 5 g yeast extract, 5 g NaCl per L) broth or on LB plates fortified with 1.5% Bacto agar at 37°C. When appropriate, antibiotics were used at the following concentrations: 5 µg/ml kanamycin, 100 µg/ml spectinomycin, 5 µg/ml chloramphenicol, and 10 µg/ml tetracycline, 1 µg/ml erythromycin with 25 µg/ml lincomycin (*mls*). Mitomycin C (MMC, DOT Scientific) was added to the medium at the indicated concentration when appropriate. Isopropyl β-D-thiogalactopyranoside (IPTG, Sigma) was added to the medium as needed at the indicated concentration.

### Strain Construction

All constructs were first introduced into the domesticated strain PY79 or into the pBS32 cured strain (DS2569) by natural competence and then transferred into the 3610-background using SPP1-mediated generalized phage transduction (18).Strains were also produced by transforming directly into the competent derivatives of 3610: DK607 (Δ*coml*) or DK1042 (Q. to L change at position 12 encoded by *comI*. All strains used in this study are listed in Table 1. All plasmids used in this study are listed in Supplemental Table S1. All primers used in this study are listed in Supplemental Table S2.

**Table 1:**
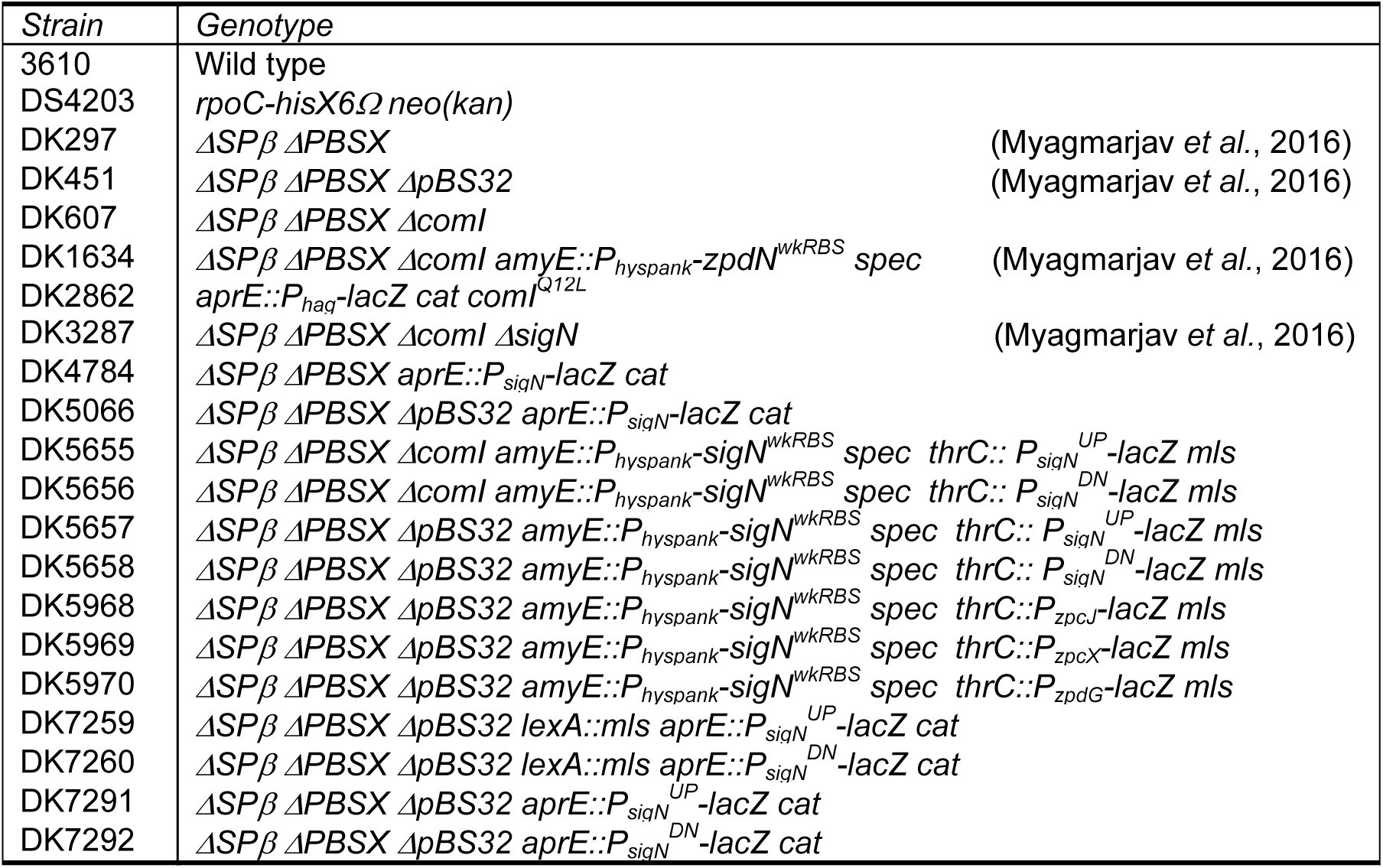
Strains.

#### *LacZ* Reporter Constructs

To generate the β-galactosidase (*lacZ*) reporter construct *aprE::P_sigN_-lacZ cat*, PCR was utilized to amplify the promoter region of *sigN* using the primer set 4500/4528 from *B. subtilis* 3610 chromosomal DNA and primer set 4438/4501 was used to amplify the *aprE* up region and *cat^R^* from DK2862 while primer set 4527/4441 was used to amplify the *aprE* down region and *lacZ* from DK2862. These DNA fragments were ligated together in a Gibson ITA assembly reaction (see below) for 1 hr at 60°C. Cementing PCR was performed using primer set 4438/441 and cleaned up using a QIAquick PCR Purification Kit (Qiagen) and transformed into DK1042.

To generate the *P_sigN_^UP^* and *P_sigN_^DN^* β-galactosidase reporter constructs at *thrC*, the promoter region of *sigN* was amplified via PCR with the primer set 6089/6090 for *P_sigN_^UP^* and 6087/6088 for *P_sigN_^DN^* from *B. subtilis* 3610 chromosomal DNA. Each PCR product was digested with *Eco*RI and *Bam*HI and cloned independently into the *Eco*RI and *Bam*HI sites of plasmid pDG1663, which carries an erythmomycin-resistance marker and a polylinker upstream of the *lacZ* gene between the two arms of the *thrC* gene to create pATB9 and pATB10 resepctively. These plasmids were transformed into DK1042.

To generate the *P_sigN_^UP^* and *P_sigN_^DN^* β-galactosidase reporter constructs at *aprE*, the first half of the promoter was PCR amplified using primers 4500 and 4707 from *B. subtilis* 3610 chromosomal DNA. The second half of the promoter was amplified using primer set 4708/4528 from *B. subtilis* 3610 chromosomal DNA. For *P_sigN_^UP^*, the flanking regions of AprE were amplified as described above. For *P_sigN_^DN^*, the flanking regions were amplified with the following primer sets: *aprE* up region and *cat^R^* (4438/4498) *aprE* down region and *lacZ* (4527/4441) from DK2862. Each promoter region was fused to the respective flanking arms of the *aprE* region using Gibson ITA assembly as described above. Fused and amplified fragments were transformed into DK1042.

To generate the *P_zpcJ_*, *P_zpcX_*, and *P_zpdG_* β-galactosidase reporter constructs, primer sets were used in the following order to amplify each promoter region: 6276/6277 (*P_zpcJ_*), 6278/6279 (*P_zpcX_*), and 6280/6281 (*P_zpdG_*). Each promoter region was digested with *Eco*RI and *Bam*HI and subsequently cloned independently into the *Eco*RI and *Bam*HI sites of plasmid pDG1663, which carries an erythmomycin-resistance marker and a polylinker upstream of the *lacZ* gene between the two arms of the *thrC* gene to create pATB12, pATB13, and pATB14 resepctively. These plasmids were transformed into DK1042.

#### lexA::mls

The *lexA::mls* insertion deletion allele was generated using a modified “Gibson” isothermal assembly protocol (19). Briefly, the region upstream of *lexA* was PCR amplified using the primer pair 5661/5662 and the region downstream of *lexA* was PCR amplified using the primer pair 5663/5664. To amplify the *erm* resistance gene, pAH52 plasmid DNA was used in a PCR reaction with the universal primers 3250/23251. Fragments were added in equimolar amounts to the Gibson ITA assembly reaction and it was performed as explained above. The completed reaction was then PCR amplified using primers 5661/5664 to amplify the assembled product. The product was transformed into DK1042.

#### Isothermal assembly reaction buffer (5X)

500 mM Tris-HCL (pH 7.5), 50 mM MgCl_2_, 50 mM DTT (Bio-Rad), 31.25 mM PEG-8000 (Fisher Scientific), 5.02 mM NAD (Sigma Aldrich), and 1 mM of each dNTP (New England BioLabs) was aliquoted and stored at −80°C. An assembly master mixture was made by combining prepared 5X isothermal assembly reaction buffer (131 mM Tris-HCl, 13.1 mM MgCl_2_, 13.1 mM DTT, 8.21 mM PEG-8000, 1.32 mM NAD, and 0.26 mM each dNTP) with Phusion DNA polymerase (New England BioLabs) (0.033 units/µL), T5 exonuclease diluted 1:5 with 5X reaction buffer (New England BioLabs) (0.01 units/µL), Taq DNA ligase (New England BioLabs) (5328 units/µL), and additional dNTPs (267 µM). The master mix was aliquoted as 15 µl and stored at −80°C.

### SPP1 Phage Transduction

To a 0.2 ml dense culture grown in TY broth (LB supplemented with 10 mM MgSO_4_ and 100 μM MnSO_4_ after autoclaving), serial dilutions of SPP1 phage stock were added. This mixture was allowed to statically incubate at 37°C for 15 minutes. A 3 ml volume of TYSA (molten TY with 0.5% agar) was added to each mixture and poured on top of fresh TY plates. The plates were incubated at 37 °C overnight. Plates on which plaques formed had the top agar harvested by scraping into a 50 ml conical tube. To release the phage, the tube was vortexed for 20 seconds and centrifuged at 5,000 × *g* for 10 minutes. The supernatant was passed through a 0.45 μm syringe filter and stored at 4 °C.

Recipient cells were grown in 2 ml of TY broth at 37 °C until stationary phase was reached. A 5 µl volume of SPP1 donor phage stock was added to 0.9 of cells and 9 ml of TY broth was added to this mixture. The transduction mixture was allowed to stand statically at room temperature for 30 minutes. After incubation, the mixture was centrifuged at 5,000 × *g* for 10 minutes, the supernatant was discarded, and the pellet was resuspended in the volume left. 100 – 200 µl of the cell suspension was plated on TY fortified with 1.5% agar, 10 mM sodium citrate, and the appropriate antibiotic for selection.

### Protein Purification

To create the SUMO-SigN fusion protein expression vector, the coding sequence of SigN was amplified from 3610 genomic DNA with primers that also introduced a *Sap*I site at the 5’ end and a *Bam*HI site at the 3’ end. This fragment was ligated into the *Sap*I and *Bam*HI sites of pTB146 to create pBM05. To purify SigN, pBM05 was expressed in Rosetta Gami II cells and grown at 37 °C until mid-log phase (∼0.5 OD_600_). IPTG was added to the cells to induce protein expression and cells were allowed to grow overnight at 16 °C. Cells were harvested by centrifugation, washed, and emulsified with EmulsiFlex-C3 (Avestin). Lysed cells were ultracentrifuged at 14,000 × g for 30 minutes at 4°C. The supernatant was mixed with Ni^2+^-NTA His•Bind resin (EMD Millipore) equilibrated with Lysis/Binding Buffer (50 mM Na_2_HPO_4_, 300 mM NaCl, 10 mM Imidazole, final pH 7.5) and allowed to incubate overnight at 4°C. The bead/lysate mixture was allowed to pack in a 1 cm separation column (Bio-Rad) and washed with Wash Buffer (50 mM Na_2_HPO_4_, 300 mM NaCl, 30 mM Imidazole, final pH 7.5). His-SUMO-SigN bound to the resin and was eluted using a stepwise elution of Wash Buffer with 50 −500 mM Imidazole and 10% glycerol to a final pH 7.5. Elutions were separated by SDS-PAGE and stained with Coomassie Brilliant Blue to verify purification. Purified His-SUMO-SigN was combined with Ubiquitin Ligase (protease) and Cleavage Buffer and incubated for at room temperature for 4 hrs to cleave the SUMO tag from the SigN protein (Butt et al., 2005 (add inREF)). The cleavage reaction was combined with Ni^2+^-NTA His•Bind resin, incubated for 1 hour at 4°C and centrifuged to pellet the resin. Supernatant was removed and dialyzed into Lysis/Binding Buffer without the Imdazole (50 mM Na_2_HPO_4_, 300 mM NaCl, 20% glycerol, final pH 7.5). Removal of the tag was confirmed by SDS-Page and staining with Coomassie Brilliant Blue.

To purify RNA polymerase, LB supplemented with kanamycin (5 µg/ml) was inoculated with an overnight culture of DK4203, which has the *rpoC-hisX6* construct integrated into the native site of *rpoC*. The cells were grown at 37 °C until they hit mid-log phase (∼0.5 OD_600_) and harvested via centrifugation. The collected cells were washed with Buffer I [10 mM Tris-HCl (pH 8.0), 0.1 M KCl, 1mM β-mercaptoethanol, 10% (v/v) glycerol] twice, resuspended in Buffer G [10 mM Tris-HCl (pH 8.0), 20% (v/v) glycerol, 10 mM imidazole, 0.5 mg/ml lysozyme], and emulsified with EmulsiFlex-C3 (Avestin). The extracts were centrifuged for 30 min at 28,000 × g twice. The supernatant was mixed with Ni^2+^-NTA His•Bind resin (EMD Millipore) equilibrated with Buffer G and allowed to go overnight at 4 °C. Collect the resin by centrifugation and wash with Buffer G. Buffer E [10 mM Tris-HCl (pH 8.0), 20% (v/v) glycerol, 500 mM imidazole] was used to elute the proteins associated with the resin and dialyzed in TGED buffer [10 mM Tris-HCl (pH 8.0), 1 mM EDTA, 0.3 mM DTT, 20% (v/v) glycerol].

To create the SUMO-LexA fusion protein expression vector, the coding sequence of LexA was amplified from 3610 genomic DNA with primers that also introduced a *Sap*I site at the 5’ end and a *Bam*HI site at the 3’ end. This fragment was ligated into the *Sap*I and *Bam*HI sites of pTB146 to create pATB11.

For the purification of LexA, pATB11 was expressed in Rosetta Gami II cells and grown at 37 °C until mid-log phase (∼0.5 OD_600_). Cells were treated the same as in the protein purification procedure for SigN (above).

### SigN Antibody Purification

One milligram of purified SigN protein was sent to Cocalico Biologicals for serial injection into a rabbit host for antibody generation. Anti-SigN serum was mixed with SigN-conjugated Affigel-10 beads and incubated overnight at 4°C. Beads were packed onto a 1 cm column (Bio-Rad) and washed with 100mM glycine (pH 2.5) to release the antibody and neutralized immediately with 2M Tris base. The antibody was verified using SDS-PAGE and stained with Coomassie Brilliant Blue Purified anti-SigN antibody was dialyzed into 1X PBS with 50% glycerol and stored at −20°C.

### Western blotting

*B. subtilis* strains were grown in LB and treated with Mitomycin C (final concentration 0.3 µg/ml) as reported in Myagmarjav, *et al* 2016. Cells were harvested by centrifugation at the different time points after treatment. Cells were resuspended to 10 OD_600_ in Lysis buffer [20 mM Tris-HCL (pH 7.0), 10 mM EDTA, 1 mg/ml lysozyme, 10 µg/ml DNAse I, 100 µg/ml RNAse I, 1 mM PMSF] and incubated for 1 hour at 37°C. 20 µl of lysate was mixed with 4 µl 6x SDS loading dye. Samples were separated by 12% sodium dodecyl sulfate-polyacrylamide gel electrophoresis (SDS-PAGE). The proteins were electroblotted onto nitrocellulose and developed with a primary antibody used at a 1:5,000 dilution of anti-SigN, 1:80,000 dilution of anti-SigA, and a 1:10,000 dilution secondary antibody (horseradish peroxidase-conjugated goat anti-rabbit immunoglobulin G). Immunoblot was developed using the Immun-Star HRP developer kit (Bio-Rad).

### β-galactosidase Assay

Biological replicates of *B. subtilis* strains were grown in LB and treated with Mitomycin C to a final concentration of 0.3 µg/ml. Cells were allowed to grow, and 1 ml was harvested by centrifugation at the different time points indicated after treatment. When IPTG (final concentration 1mM) was used, cells grew to an OD_600_ 0.6 and 1 ml was harvested. Cells were resuspended in 1 ml of Z-buffer (40 mM NaH_2_PO_4_, 60 mM Na_2_HPO_4_, 1mM MgSO_4_, 10 mM KCl, and 38 mM β-mercaptoethanol) with 0.2 mg/ml of lysozyme and incubated at 30 °C for 15 minutes. Each sample was diluted accordingly with Z-buffer to 500 µl. The reaction was started with 100 µl of 4 mg/ml O-nitrophenyl β-D-galactopyranoside (in Z buffer) and stopped with 1M Na_2_CO_3_ (250 µl). The OD_420_ of each reaction was noted and the β-galactosidase specific activity was calculated using this equation: [OD_420_/(time × OD_600_)] × dilution factor × 1000.

### Collection of cells for Rend-seq

Overnight cultures were back diluted 1:100 in LB and grown at 37 C shaking.

When the cultures reached an OD600 of 0.1 they were treated with either 1 ug/ml MMC (DK297 and DK3287) or 1mM IPTG (DK1634). The zpdN over expression strain was harvested 1 hour after induction by IPTG. Cells treated with MMC were collected after 2 hours. After treatment, 10 ml of each culture was mixed with 10 ml of ice cold methanol and spun down at 3220 ×g at 4 °C for 10 minutes. Supernatant was discarded and cell pellets were frozen in liquid nitrogen and stored at −80°C. For RNA extraction, the thawed pellets were resuspended in 1 ml of Trizol reagent (Thermo Fisher, Waltham, MA) and added to FastPrep lysis matrix B 2 ml tubes with beads (MP Biomedicals). Cells were disrupted in a Bead Ruptor 24 (Omni International, Kennesaw, GA) twice for 40 seconds at 6.0 M/s. 200 μl of chloroform was and were kept at room temperature for 2 minutes after vigorous vortexing. Mixture was spun down at 18,200 ×g 30 minutes at 4°C. The aqueous phase (∼600 μl) was precipitated with 900 μl of isopropanol for 10 minutes at room temperature. The RNA pellet was collected and washed with 80% ethanol.

### Rend-seq library preparation

RNA was prepared for Rend-seq as described in detail in Lalanne et al. 2018 and DeLoughery et al. 2018. In brief, 5-10 μg RNA was DNAse treated (Qiagen) and rRNA was depleted (MICROBExpress ThermoFisher). rRNA depleted RNA was fragmented by first heating the sample to 95°C for 2 min and adding RNA fragmentation buffer (1x, Thermo Fisher) for 30 seconds at 95°C and quenched by addition of RNA fragmentation stop buffer (ThermoFisher). RNA fragments between 20 and 40 bp were isolated by size excision from a denaturing polyacrylamide gel (15%, TBE-Urea, 65 min., 200 V, ThermoFisher). RNA fragments were dephosphorylated using T4 polynucleotide kinase (New England Biolabs, Ipswitch, MA), precipitated, and ligated to 5’ adenylated and 3’-end blocked linker 1 (IDT 5 μM) using T4 RNA ligase 2, truncated K227Q. The ligation was carried out at 25°C for 2.5 hours using <5 pmol of dephosphorylated RNA in the presence of 25% PEG 8000 (ThermoFisher). cDNA was prepared by reverse transcription of ligated RNA using Superscript III (ThermoFisher) at 50°C for 30 min. with primer oCJ485 (IDT, Coralville, Iowa) and the RNA was hydrolyzed. cDNA was isolated by PAGE size excision (10% TBE-Urea, 200V, 80 min., ThermoFisher). Single stranded cDNAs were circularized using CircLigase (Illumina, San Diego, CA) at 60°C for 2 hours. Circularized cDNA was the template for PCR amplification using Phusion DNA polymerase (New England Biolabs) with Illumina sequencing primers, primer o231 (IDT) and barcoded indexing primers (IDT). After 6 – 10 rounds of PCR amplification, the product was selected by size from a non-denaturing PAGE (8% TB, 45 min., 180V, Life Technologies. For dataset names and barcode information see Table S3.

### RNA-sequencing and data analysis

Sequencing was performed on an Illumina HiSeq 2000. The 3’ linker sequences were stripped. Bowtie v. 1.2.1.1 (options -v 1 -k 1) was used for sequence alignment to the reference genome NC 000964.3 (*B. subtilis* chromosome) and KF365913.1 (*B. subtilis* plasmid pBS32) obtained from NCBI Reference Sequence Bank. To deal with non-template addition during reverse transcription, reads with a mismatch at their 5’ end had their 5’ end re-assigned to the immediate next downstream position. The 5’ and 3’ ends of mapped reads between 15 and 45 nt in sizes were counted separately at genomic positions to produce wig files. The wig files were normalized per million non-rRNA and non-tRNA reads for each sample. Shadows were removed from wig files first by identifying the position of peaks and then by reducing the other end of the aligned reads by the peak’s enrichment factor to produce the final normalized and shadow removed wig files. Gene regions were plotted in MATlab.

### Electromobility Shift Assays

To perform electromobility shift assays, LexA was purified from *E. coli* as outlined above. The control promoter, *P_recA_*, was amplified using the primer set 6252/6253, *P_sigN_^UP^* was amplified using the primer set 6089/6090, *P ^DN^* was amplified using the primer set 6087/6088, and *P_sigN_^UP^*\* (LexA site scrambled) was amplified using the primer set 6089/6284 and 6090/6283 from *B. subtilis* 3610 genomic DNA. The *P_sigN_^UP^*\* fragments were ligated using Gibson ITA assembly as outlined above. All fragments were cleaned up using the QIAquick PCR Purification Kit (Qiagen). Each DNA probe was end labeled with γ^32^P-ATP with T4 PNK (New England Biolabs). Excess nucleotide was removed using G-50 microcolumns (GE Life Technologies). DNA binding reactions contained 4 nM of the DNA probe and a specific concentration of purified LexA protein (either 1, 5, 10, 50, 100, or 500 nM). Reactions were carried out in binding buffer (100 mM HEPES pH 7.5, 100 mM Tris-HCl, 50% glycerol, 500 mM NaCl, 10 mM EDTA, 10 mM DTT) supplemented with 100 µg/ml bovine serum albumin (BSA) and 10 ng/µl poly(dI-dC). All reactions were incubated for 45 minutes at room temperature. Protein-DNA complexes were resolved on a 6% TGE polyacrylamide gel. Gels were dried at 80°C for 90 minutes and exposed to a storage phosphor screen overnight. Gels were imaged with a Typhoon 9500 (GE Life Sciences).

### *in vitro* Transcription

DNA template (50 ng) was mixed with either RNAP only (250 nM) or with RNAP plus SigN (1000 nM) per reaction. Each reaction was incubated for 15 minutes at 37 °C in 25 µl total reaction volume including the transcription buffer [18mM Tris HCl (pH 8.0), 10 mM MgCl_2_, 30 mM NaCl, 1mM DTT, 250 µM GTP, 100 µM ATP, 100 µM CTP, 5 µM UTP, and ∼2 µCi [α-^32^P] UTP] to produce multiple round transcription. To stop the reaction, 25 µl of 2X Stop/Gel Loading solution (7M urea, 10 mM EDTA, 1% SDS, 2X TBE, 0.05% bromophenol blue) was used. Samples were ran on a 5% denaturing acrylamide gel [5% Acrylamide (19:1 acryl:bis), 7M urea, 1X TBE] for 3 hours at 200V. Gels were imaged with a Typhoon 9500 (GE Life Sciences).

### Data and software availability

Ribosome profiling and RNA-sequencing are available at the Gene Expression Omnibus under accession number GSEXXXXX, which can be accessed using the reviewer token mvcjogkcvxglfez. Data were analyzed using custom Matlab scripts which are available upon request. Note to editor and reviewers: Rend-seq analysis has been submitted for database access, accession number is pending and will be provided upon revision.

## Results

### SigN is repressed by LexA

SigN (formerly ZpdN) is a sigma factor homolog encoded on the plasmid pBS32 that is necessary and sufficient for pBS32-mediated cell death (17). Consistent with previous results, treatment of cells deleted for the PBSX and SPβ prophages (14, 20–23) with the DNA damaging agent mitomycin C (MMC) caused a 3-fold decrease in optical density (OD) from peak absorbance, and the decrease in OD was abolished in cells also deleted for *sigN* (17) (Fig 1A). To determine the effect of MMC on cell viability, viable counting was performed by dilution plating over a time-course following MMC addition. Addition of MMC caused a rapid and immediate decline in colony forming units such that the number of viable cells decreased three-orders of magnitude even as the OD increased for three doublings (compare Figs 1A and 1B). As with loss of OD, mutation of *sigN* abolished the MMC-dependent decrease in cell viability (Fig 1B). We conclude that pBS32-mediated cell death occurs prior to, and independent of, transient cell growth and the subsequent decline in OD, and that SigN is required for all pBS32-dependent death-related phenotypes thus far observed.

**Figure 1:**
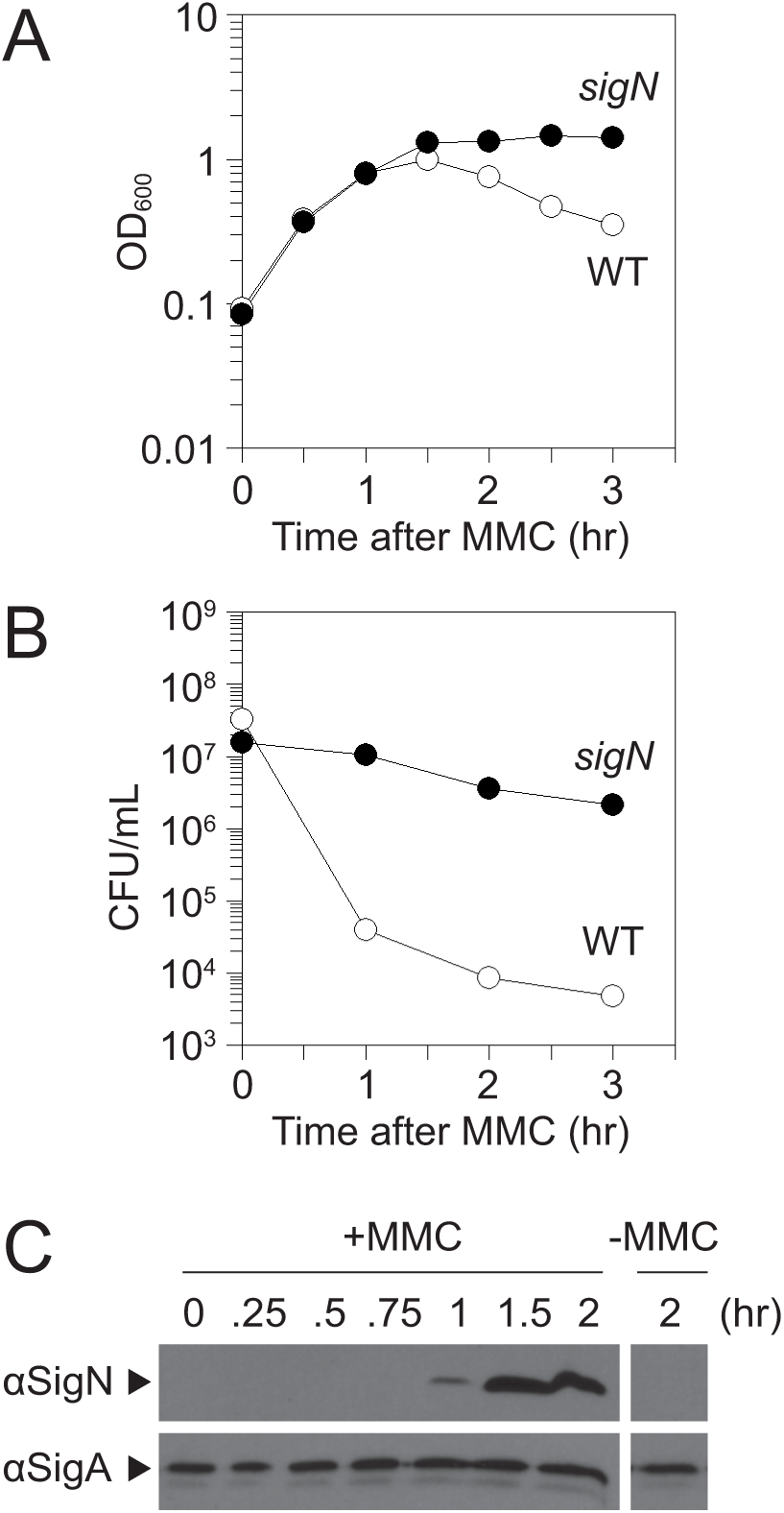
SigN is required for loss of cell viability after MMC treatment. A) Optical density (OD_600_) growth curve of wild type (open circles, DK607) and sigN mutant (closed circles, DK3287). X-axis is time of spectrophotometry after MMC addition. B) Colony forming unit growth curve of wild type (open circles, DK607) and sigN mutant (closed circles, DK3287). X-axis is time of dilution plating after MMC addition. C) Western blot analysis of wild type DK607 cell lysates harvested at the indicated time after MMC addition and probed with either anti-SigN antibody or anti-SigA antibody. On right is a single panel of the same strain for comparison 2 hours after mock MMC addition.

To determine if and when SigN was expressed relative to MMC treatment, Western blot analysis was conducted. SigN protein was first detected one hour after MMC treatment and continued to increase in abundance thereafter, whereas the vegetative sigma factor, SigA (σ^A^), was constitutive and did not increase (Fig 1C). We noted that loss of cell viability appeared to occur soon after MMC addition, perhaps prior to observable SigN protein (e.g. 0.5 hrs. after addition, Fig 1B), and thus we inferred that SigN was expressed and active at levels below the limit of protein detection. To determine whether SigN expression occurs soon after MMC treatment, the upstream intergenic region of *sigN* (P*_sigN_*) (Fig 2A) was cloned upstream of the gene encoding β-galactosidase, *lacZ*, and inserted at an ectopic site in the chromosome (*aprE::P_sigN_-lacZ*). Expression from P*_sigN_* was low but increased 10-fold within an hour after MMC addition (T_1_), and the increase in expression was not dependent on the presence of pBS32 (Fig 3A).

**Figure 2:**
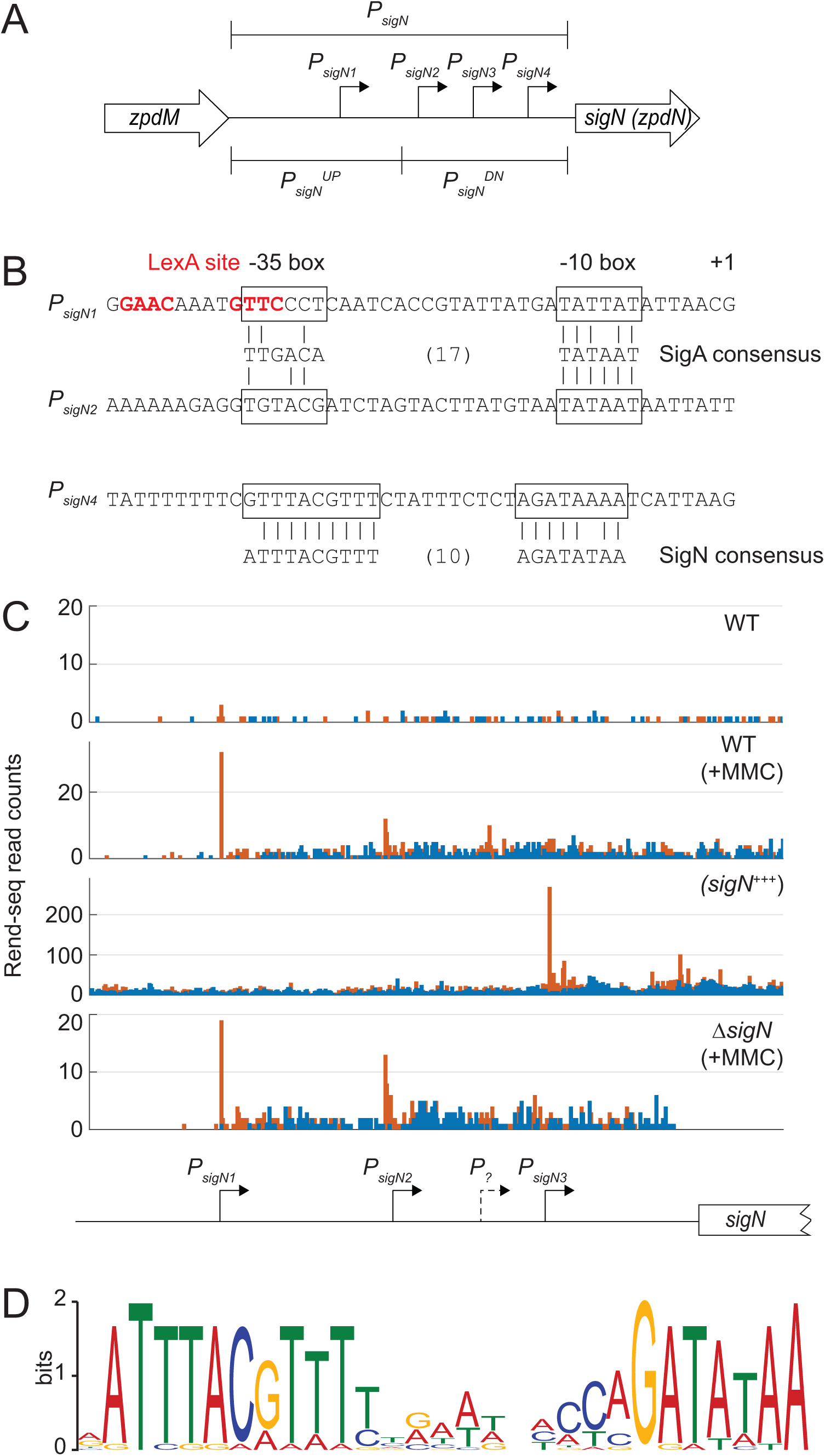
SigN promoter region. (A) A schematic of the promoter region of *sigN*. Open arrows indicate reading frames. Bent arrows indicate promoters. Promoter regions are indicated by brackets. (B) Promoter sequences. Boxes surround −35 and −10 regions relative to the +1 transciptional start site. Below the promoters are SigA and SigN consensus sequences with vertical lines to indicate a consensus match. C) REND-seq data for the indicated genotypes: WT (DK607), WT+MMC (DK607 induced for 1 hr with MMC), *sigN*^+++^ (DK1634 induced for 1 hr with 1 mM IPTG), and *ΔsigN*+MMC (DK3287 induced for 1 hr with MMC). Orange peaks represent 5’ ends and blue peaks represent 3’ ends. Below is a cartoon indicating the location of the promoter believed to be responsible for transcriptional start sites predicted above relative to the *sigN* coding region. Note, the peaks stop abruptly in the last panel due to deletion of the *sigN* gene. Information on RENDseq is included in Table S3. (D) SigN consensus sequence generated by MEME sequence analysis using the promoters listed in Table 2.

**Figure 3:**
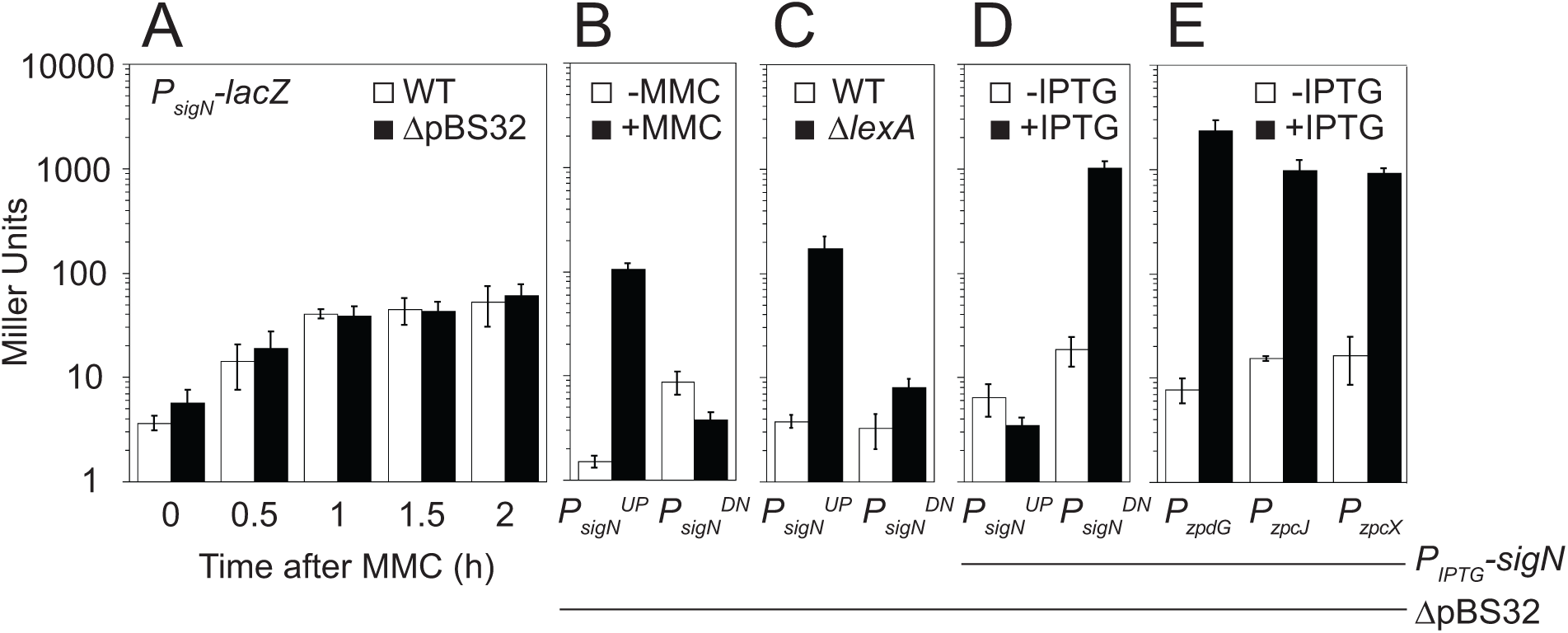
The *sigN* promoter region is repressed by LexA and autoactivated. **(A)** β-galactosidase activity of a *P_sigN_-lacZ* reporter in the presence (open bars) and absence (closed bars) of pBS32 measured at the indicated timepoints following 800nM MMC addition. The following strains were used to generate this panel: DK4784 (WT) and DK5066 (Δ*pBS32*). B) β-galactosidase activity of either a *P_sigN_^UP^-lacZ* or *P_sigN_^DN^-lacZ* reporter in the presence (closed bars) and absence (open bars) of 800nM MMC (1 hour incubation). The following strains were used to generate this panel: DK5657 (*P_sigN_^UP^-lacZ* Δ*pBS32*) and DK5658 (*P_sigN_^DN^-lacZ* Δ*pBS32*). C) β-galactosidase activity of either a *P_sigN_^UP^-lacZ* or *P_sigN_^DN^-lacZ* reporter i565n the presence (closed bars) and absence (open bars) of LexA. The following strains were used to generate this panel: DK7291 (*P_sigN_^UP^-lacZ* Δ*pBS32*), DK7292 (*P_sigN_^DN^-lacZ* Δ*pBS32*), DK7259 *P_sigN_^UP^-lacZ* Δ*pBS32 lexA*), and DK7260 (*P_sigN_^DN^-lacZ* Δ*pBS32 lexA*). D) β-galactosidase activity of either a *P_sigN_^UP^-lacZ* or *P_sigN_^DN^-lacZ* reporter in strain containing and IPTG-inducible SigN construct grown in the presence (closed bars) and absence (open bars) of 1 mM IPTG. The following strains were used to generate this panel: DK5657 (*P_sigN_^UP^-lacZ* Δ*pBS32*) and DK5658 (*P_sigN_^DN^-lacZ* Δ*pBS32*). E) β-galactosidase activity of a *P_zpdG_-lacZ, P_zpcJ_-lacZ* or *P_zpcX_-lacZ* reporter in strain containing and IPTG-inducible SigN construct grown in the presence (closed bars) and absence (open bars) of 1 mM IPTG. The following strains were used to generate this panel: DK5970 (*P_zpdG_-lacZ* Δ*pBS32*), DK5968 (*P_zpcJ_-lacZ* Δ*pBS32*), and DK5969 (*P_zpcX_-lacZ* Δ*pBS32*). Error bars are the standard deviation of three replicates. Data used to generate each panel is included in Table S4-S8.

**Table 2:**
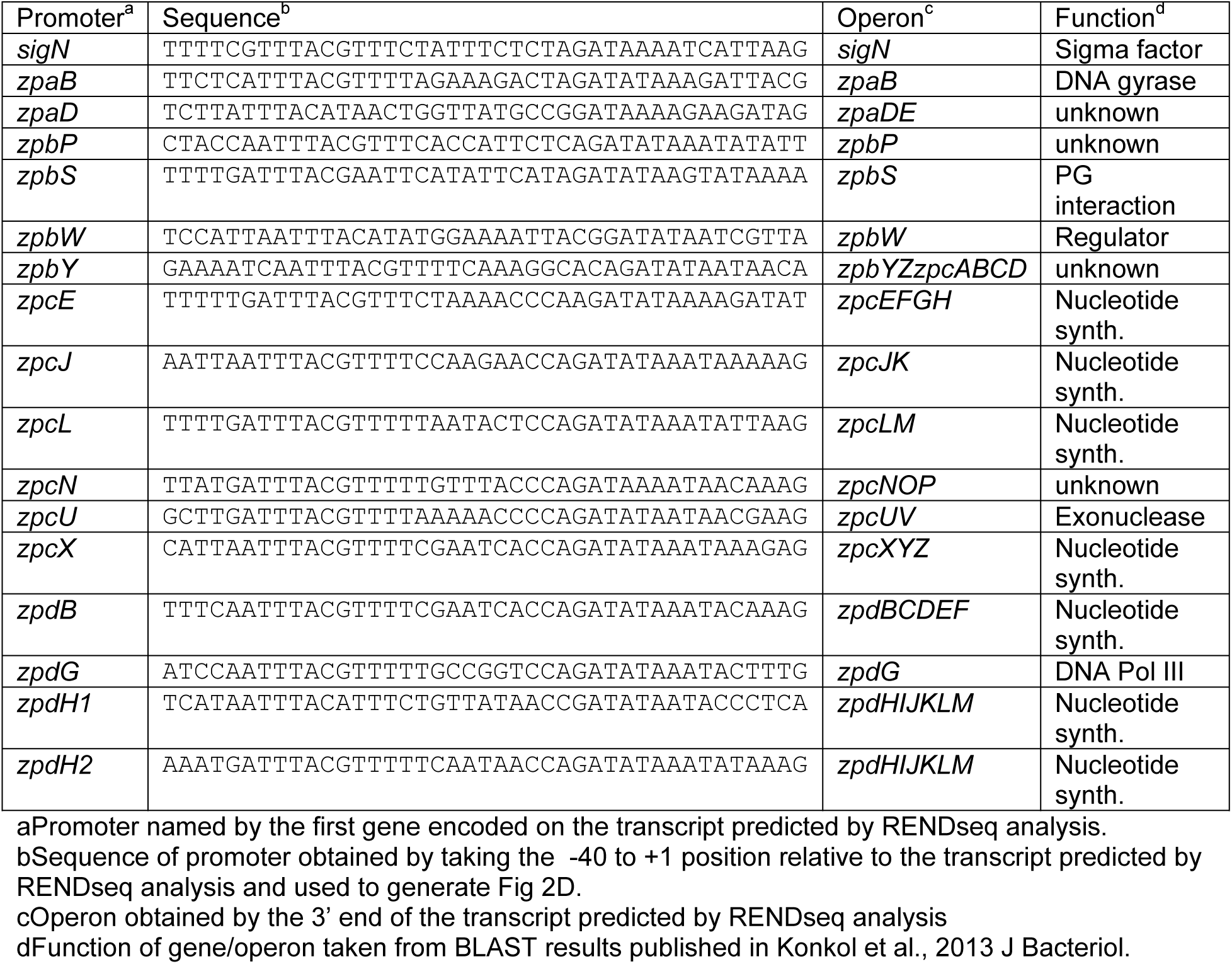
SigN-dependent promoters on pBS32.

To map the MMC-response within the *sigN* promoter region, we split the P*_sigN_* region into two fragments, an upstream fragment called *P_sigN_^UP^* and a downstream fragment called *P_sigN_^DN^* (Fig 2A). Both fragments were cloned upstream of *lacZ* and separately integrated into an ectopic site of the chromosome in a strain deleted for pBS32 and both chromosomal prophages, PBSX and SPβ. Basal expression from P*_sigN_^UP^* was at background levels but increased 100-fold when MMC was added (Fig 3B). In contrast, expression from P*_sigN_^DN^* was expressed at a constitutively low level and did not increase upon addition of MMC (Fig. 3B). We conclude that transcription of *sigN* is activated by MMC treatment, that the P*_sigN_^UP^* region contains an MMC-responsive promoter, and that MMC-dependent expression was controlled by a chromosomally-encoded regulator as induction was not dependent on the presence of pBS32.

One candidate for an MMC-responsive, chromosomally-encoded regulator is the transcriptional repressor protein LexA. LexA often binds to sequences that overlaps promoters to inhibit access of RNA polymerase holoenzyme (24, 25), and sequence analysis predicted a putative LexA-inverted repeat binding site located within the P*_sigN_^UP^* fragment (26, 27) (Supp. Fig 1). Moreover, target promoters are exposed and expression is de-repressed when LexA undergoes auto-proteolysis upon DNA damage like that caused by MMC (24, 25, 28). To determine if P*_sigN_^UP^* was LexA repressed, LexA was mutated in a background deleted for pBS32 and the two chromosomal prophages, PBSX and SPβ. Mutation of *lexA* dramatically increased expression from P*_sigN_^UP^* but not P*_sigN_^DN^* (Fig 3C). We conclude that LexA either directly or indirectly inhibits expression of a promoter present in P*_sigN_^UP^*.

One way that LexA might inhibit expression from P*_sigN_^UP^* is if it bound directly to the DNA. To determine whether LexA bound directly to the P*_sigN_^UP^* region, LexA was purified and added to various labeled DNA fragments in an electrophoretic mobility shift assay (EMSA). Consistent with direct, high-affinity binding, purified LexA caused an electrophoretic mobility shift in both the previously established target promoter P*_recA_* (25) (Fig 4A) and the P*_sigN_^UP^* promoter region (Fig 4B) at protein levels as low as 1 nM. LexA binding was specific as the affinity was reduced 500-fold for the P*_sigN_^DN^* promoter (Fig 4C). Moreover, LexA binding was specific for the putative LexA inverted repeat sequence as mutation of the sequence (GAAC > <u>TT</u>AC) within P*_sigN_^UP^* reduced binding affinity 100-fold (Fig 4D). We conclude that LexA binds to the P*_sigN_^UP^* promoter region and represses transcription.

**Figure 4:**
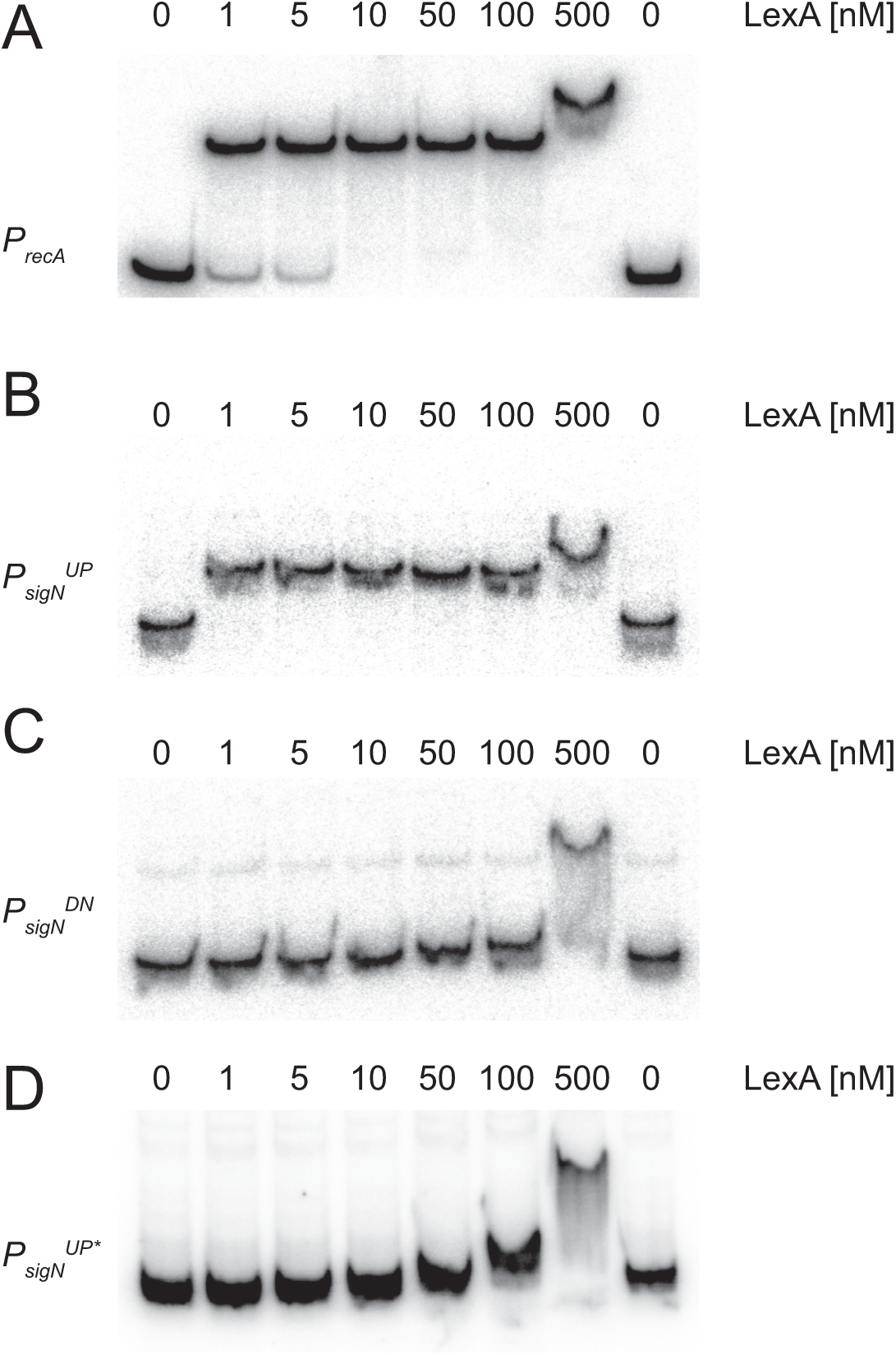
LexA binds to the *P_sigN_^UP^* promoter region. Electrophoretic mobility shift assays were performed with radiolabeled DNA of PrecA (A), PsigNUP (B), PsigNDN (C) and PsigNUP* mutated for the putative LexA binding site (D). Purified LexA protein was added to each reaction at the indicated concentration.

LexA often binds overtop of promoter elements (16), and sequence analysis suggested that the LexA inverted repeat in P*_sigN_^UP^* might rest immediately upstream of, an overlap with, a putative SigA-dependent −35 promoter element (Fig 2B). To determine whether P*_sigN_^UP^* contained a SigA-dependent promoter, RNA polymerase (RNAP) holoenzyme with SigA bound was purified from *B. subtilis* and used in an *in vitro* transcription reaction (Fig 5). Consistent with promoter activity, transcription product was observed when SigA-RNAP was mixed with either a known SigA-dependent promoter control *P_veg_* (Fig 5A, left lane), or the experimental P*_sigN_^UP^* (Fig 5B, left lane). A transcription product was also observed when SigA-RNAP was mixed with the P*_sigN_^DN^* promoter fragment (Fig 5C, left lane), consistent with low level constitutive expression observed from reporters with that fragment (Fig 3B). We conclude that there are two SigA-dependent promoters within the *P_sigN_* region, one within the P*_sigN_^UP^* fragment and one within P*_sigN_^DN^* fragment.

**Figure 5:**
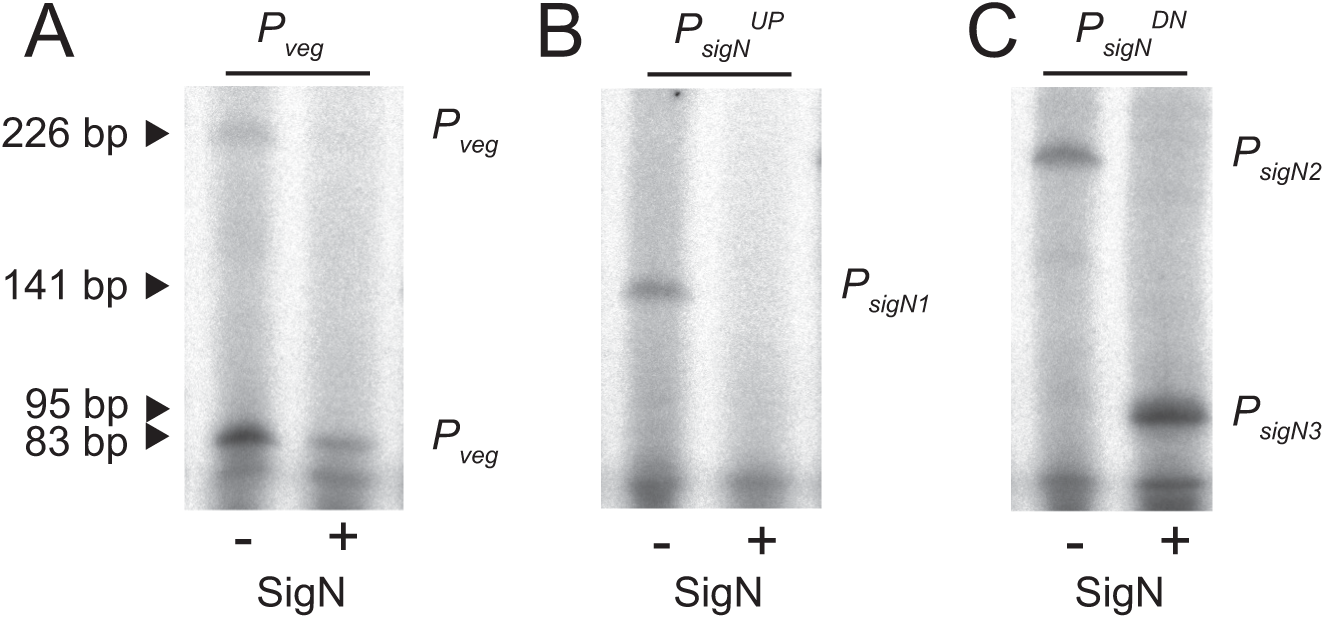
SigN is a sigma factor that drives transcription *in vitro*. *In vitro* transcription assays using *P_veg_* (left), *P_sigN_^UP^* (middle), and *P_sigN_^DN^* (right) promoter fragments in the presence (+) and absence (-) of 5X molar ratio of SigN added to RNA polymerase holoenzyme purified from *B. subtilis*. The predicted transcriptional products resulting from *P_sigN1_*, *P_sigN2_*, and *P_sigN4_* are indicated. Two products were observed from *P_veg_* likely due to proper termination (short product) and terminator read-through (long product).

To determine transcriptional start sites, Rend-seq (end-enriched RNA-seq) analysis was performed for the entire *B. subtilis* transcriptome in the presence and absence of MMC-treatment (Fig 2B). Rend-seq achieves end-enrichment by sparse fragmentation of extracted RNAs, which generates fragments containing original 5’ and 3’ ends, as well as a lower amount of fragments containing internal ends (29, 30). Rend-seq indicated that expression of sign was low in the absence of induction (Fig 2C) but a 5’ end appeared within the P*_sigN_^UP^* region when MMC was added, the location of which was consistent with the SigA −10 promoter element predicted earlier (Fig 2B) and supported later by *in vitro* transcription (Fig 5A, left lane). We define the SigA-dependent promoter within P*_sigN_^UP^* as *P_sigN1_*. Rend-seq also indicated a weak but MMC-independent 5’ end within P*_sigN_^DN^* that was consistent with the *in vitro* transcription product originating from that fragment (Fig 5B, left lane). Moreover, sequences consistent with −35 and −10 promoter elements were identified upstream of the 5’ end within P*_sigN_^DN^* (Fig 2B). We define the weak constitutive SigA-dependent promoter within P*_sigN_^DN^* as *P_sigN2_*. We conclude that there are two SigA-dependent promoters driving *sigN* expression, and that *P_sigN1_* is both strong and LexA-repressed.

### SigN is a sigma factor that activates its own expression

Rend-seq analysis also indicated a second 5’ end within P*_sigN_^DN^* fragment that would result in a slightly shorter transcript (Fig 2B, peak marked *P_?_*). The shorter transcript could indicate either a highly specific RNA cleavage site in the 5’ upstream untranslated region of *sigN*, or the presence of a third promoter with an individual start site. If there was a second promoter within P*_sigN_^DN^*, the promoter was presumably not dependent on SigA as only one SigA-dependent transcript was observed from this fragment in *in vitro* transcription assays (Fig 5B). One candidate for an alternative sigma factor that could drive expression of the third putative promoter is SigN itself. SigN is homologous to extra-cytoplasmic function (ECF) sigma factors and ECF sigma factors are often autoregulatory (31). Consistent with autoactivation, induction of SigN increased expression from P*_sigN_^DN^*-lacZ 100-fold but did not increase expression from P*_sigN_^UP^* (Fig 3C). We conclude that *sigN* expression is controlled by at least three promoters: a LexA-repressed SigA-dependent promoter P*_sigN1_*, a weak constitutive SigA-dependent promoter *P_sigN2_*, and a third promoter that was SigN-dependent.

One way in which a promoter could be SigN-dependent is if SigN is a *bona fide* sigma factor that directs its transcription. To determine whether SigN had sigma factor activity, RNAP-SigA holoenzyme was purified from *B. subtilis* and purified SigN protein was added in 5-fold excess in *in vitro* transcription reactions(32–34). Addition of SigN reduced levels of the SigA-dependent *P_veg_*, *P_sigN1_*, and the *P_sigN2_*-derived transcripts, consistent with SigN competing with, and displacing, SigA from the RNA polymerase core (Fig. 5, right lanes). Moreover, a new shorter transcript appeared within *P_sigN_^DN^* that was SigN-dependent (Fig 5C, right lane). To map the location of the shorter transcript, Rend-seq was conducted on a strain that was artificially induced for SigN expression. Consistent with the *in vitro* transcription results, an intense SigN-dependent 5’ end was detected within the *P_sigN_^DN^* region which we infer is due to the presence of a promoter here called, *P_sigN3_* (Fig 2C). We note that the *P_sigN3_*-dependent transcript did not align with the original transcript peak from *P_?_* indicated by Rend-seq analysis and thus at least three and possibly more promoters may be present upstream of *sigN*. Moreover, both the *P_?_* and P_sigN3_-dependent peaks in the MMC-treated REND-seq, were abolished in *sigN* mutant cells (Fig 2C). Nonetheless, we conclude that SigN is a bona fide sigma factor that is necessary and sufficient for inducing expression from *P_sigN3_*.

Mapping of the Rend-seq transcriptional start site, allowed prediction of the *P_sigN3_* promoter sequence (Fig 2B). To determine a SigN consensus sequence, 40 base pairs of sequence upstream of each pBS32 5’ end of transcript as determined by Rend-seq analysis after SigN artificial expression were collected and compiled by MEME (35) (Fig 2D). A consensus sequence emerged that was consistent with the −35 and −10 regions predicted by distance analysis for *P_sigN3_* (Fig 2B). Three separate promoter regions predicted thought to be regulated by SigN were cloned upstream of a promoter-less *lacZ* gene and inserted at an ectopic site in the chromosome in a strain deleted for pBS32. In each case, the expression of the reporter was low during normal growth conditions but increased 100-fold when *sigN* was induced with IPTG (Fig 3E). We conclude that SigN is a plasmid-encoded sigma factor that is necessary and sufficient for the expression of a regulon genes encoded on pBS32, and we infer that the expression of one or more genes within the SigN regulon is responsible for pBS32-mediated cell death.

## DISCUSSION

Previously published results showed that the SigN primary sequence exhibited homology to other well-known sigma factors present in *B. subtilis* (17) and here we show that SigN exhibits sigma factor activity *in vitro*. Moreover, using Rend-seq analysis we determine the regulon of genes under SigN control and use transcriptional start sites and to identify a consensus binding sequence (Fig 2C). While plasmid-encoded sigma factors are rare, SigN is conserved on *Bacillus* plasmids that are closely related to pBS32 such as pLS32 or pBUYP1. Alternative sigma factors or analogs thereof are sometimes encoded within prophage elements (36–39), and pBS32 encodes what appears to be a cryptic prophage. Whether pBS32 in its entirety is a P1-like plasmid prophage (40),or whether a phage secondarily lysogenized into a preexisting plasmid is unknown but pBS32 in its entirety appears to be released on cell death in a capsid-dependent DNase-resistant form (17). Regardless, induction of SigN is necessary and sufficient to cause cell death in a manner dependent on pBS32.

Similar to, and perhaps consistent with, other lysogenic prophages in *B. subtilis*, DNA damage caused by MMC triggers hyper-replication of pBS32 and initiates pBS32-mediated cell death (16, 17, 41). Here we show that MMC induces the plasmid via the chromosomally-encoded transcriptional repressor LexA (Fig 6). LexA tightly represses the *P_sigN1_*promoter, and MMC-mediated DNA damage promotes the auto-proteolysis of LexA (28, 42–44). De-repression of *P_sigN1_* leads to high-level *sigN* expression and once expressed, SigN locks the system into an activated state by positive feedback at the P*_sigN3_* promoter. SigN directs not only its own expression but an entire regulon on pBS32 which includes many genes homologous to those involved in nucleotide metabolism and DNA replication (Table 2). Thus, SigN activation cause pBS32 copy number to increase 100-fold and either directly or indirectly promote cell death.

**Figure 6.**
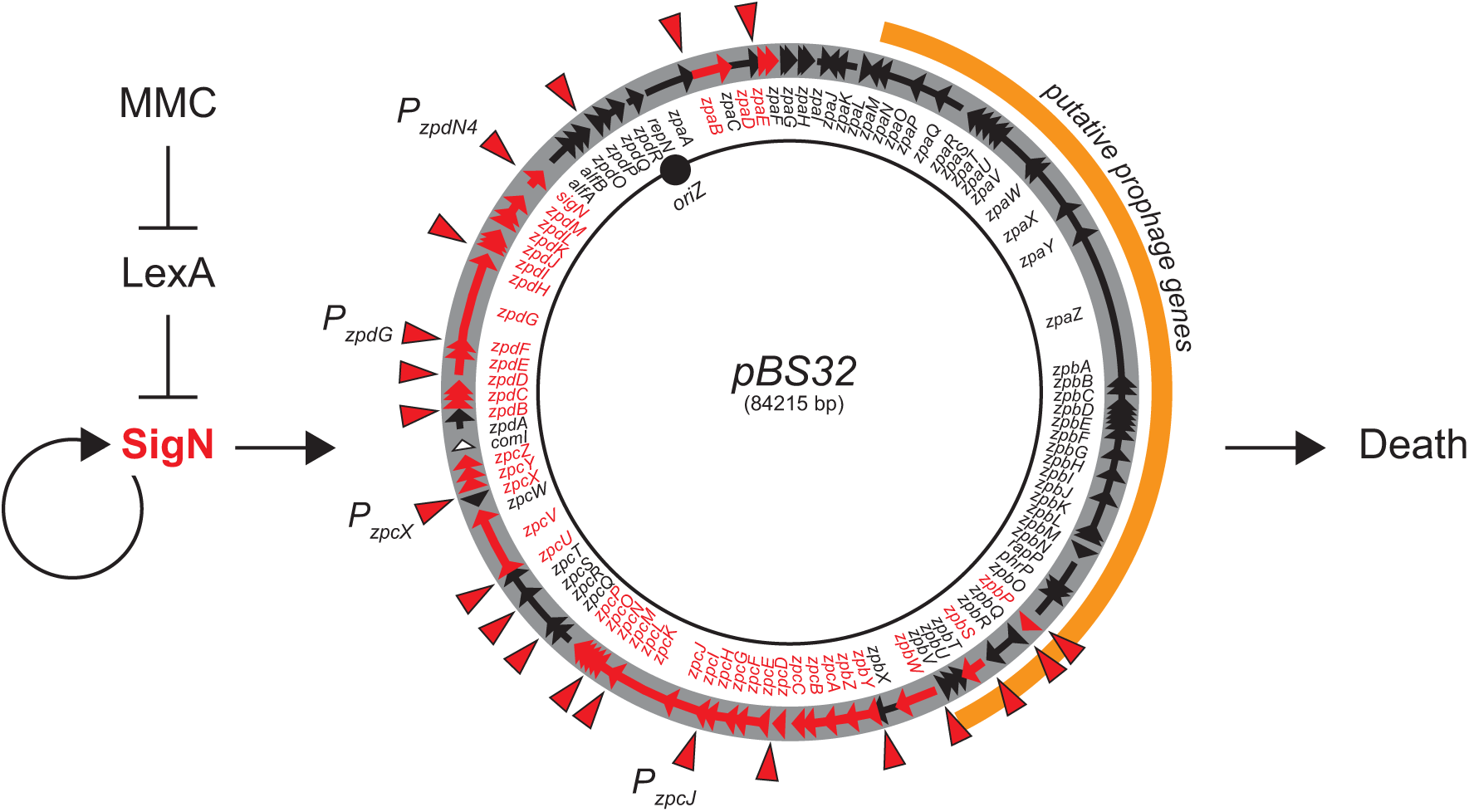
Model of pBS32-mediated cell death. MMC-mediated DNA damage causes LexA autoproteolysis and derepression of sigN expression. SigN is a sigma factor that directs RNA polymerase to increase its own expression (creating positive feedback) and the expression of a regulon of genes on pBS32. Activation of genes within the SigN-regulon results in cell death. pBS32 represented as a circle. Arrows within the circle indicate reading frames. Reading frames and gene names that are expressed by SigN are indicated in red. The location of SigN-dependent promoters is indicated by red carets. T bars indicate inhibition and arrows indicate activation.

How pBS32 actually kills cells is still unknown. Here we show that cells activated for SigN in the presence of pBS32 continue to grow for three generations even as cell viability rapidly declines. Thus, toxicity likely isn’t due to direct inhibition of essential components and instead, something essential is depleted, diluted through growth and not replaced. Hyper-replication of the plasmid may deplete nucleotide pools but at present we cannot determine whether hyper-replication and death are linked or separate phenotypes. Finally, death might be mediated by the prophage structural and lytic genes, but we note that SigN-dependent promoters appear to be largely excluded from the prophage region and while prophage gene expression increases, the increase may be largely due to the increase in plasmid copy number. Finally, large deletions of the prophage structural genes were insufficient to abolish pBS32-mediated cell death (17). Thus, prophage gene expression may be separate from SigN-mediate death as well.

Members of the extracytoplasmic sigma factor (ECF) family are typically induced by extracellular signals and promote gene expression to adapt to environment stress (45, 46). Here we show that SigN is a functional ECF-like sigma that responds to internal signals in the form of DNA damage and in turn, promotes cell death. Why cells encode a sigma factor that induces cell death is unknown. Moreover, SigN appears to be unlike most ECF sigma factors as it does not appear to be regulated by a co-expressed cognate anti-sigma. Thus, if and how SigN is regulated independently of the DNA damage response, is unknown. We note however that there is a third weak but constitutive promoter *P_sigN2_*that also drives expression of SigN. The function of *P_sigN2_* and why *P_sigN2_* is insufficient to promote SigN-mediated cell death is unknown. We speculate however, that *P_sigN2_* may either provide for additional environmental regulation on SigN or be an irrelevant vestige of former regulation. Ultimately, why *B. subtilis* retains a potentially lethal plasmid and a sigma factor that promotes cell death is unknown.

## ACKNOWLEDGEMENTS

Alyssa Ball, Ryan Chaparian, Felix Dempwolff, Masaya Fujita, Kate Hummels, Bat-Erdene Magyarjav, Reid Oshiro, Sundharraman Subramanian, Lauren Wahle and Julia van Kessel for technical support. This work was supported by NIH R35GM124732 to GWL and NIH Grant R35GM131783 to DBK.

